# The Impact of Lethal Recessive Alleles on Bottlenecks with Implications for Conservation Genetics

**DOI:** 10.1101/089151

**Authors:** R. B. Campbell

## Abstract

When a bottleneck occurs, lethal recessive alleles from the ancestral population provide a genetic load. The purging of lethal recessive mutations may prolong the bottleneck, or even cause the population to become extinct. But the purging is of short duration, it will be over before near neutral deleterious alleles accumulate. Lethal recessive alleles from the parental population and near neutral deleterious mutations which occur during a bottleneck are temporally separated threats to the survival of a population. Breeding individuals from a large population into a small endangered population will provide the benefit of viable alleles to replace near neutral deleterious alleles but also the cost of lethal recessive mutations from the large population.

## 1. Introduction

Bottlenecks are a foundation of evolution because the small population size facilitates genetic drift. One aspect is that small population size allows the fixation of near neutral deleterious mutations (DM) which can cause the extinction of a population. Ballick et al. (2015) have noted that relaxed selection (perhaps density dependent selection including a form of logistic selection) during a bottleneck may enhance the effect of decreased population size causing deleterious mutations to become near neutral and allowing them to increase due to mutation pressure. The accumulation of DM explains the disappearance of Neanderthals, with limited introgression of their genes into the human genome (Harris and Nielsen, 2016). These DM are not lethal and are seldom recessive. Lethal recessive mutations (LR) are not fixed in small populations because they are lethal, but the purging of LR from the larger ancestral population may prolong or exacerbate a bottleneck, perhaps resulting in extinction of the bottlenecked population. The purging of LR occurs on a much shorter time scale than the fixation of DM, hence provides an alternative mechanism to DM for the extinction of a population which passes through a bottleneck rather than a means to facilitate the accumulation of DM.

This paper considers the affect of LR from the ancestral population on bottlenecks. The equilibrium number of LR per individual in a population increases with population size, hence the genetic load from lethal recessive mutations when a bottleneck occurs will be greater than the equilibrium level for the size of the bottleneck. Our analysis employs estimates for the size of the human population, size of the human genome, and frequency of LR in human populations which are taken from the literature.

## 2. The number of lethal recessive mutations

There are two formulae in Crow and Kimura (1970) which provide the frequency *x* of LR in a population. On page 259 is the formula

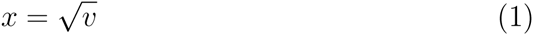

where *x* is the relative frequency of LR at a locus and *v* is the mutation rate
to LR at that locus. On page 447 is the formula

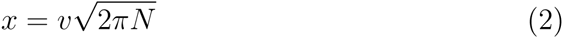

(we are considering a population of *N* diploid, hence 2*N* haploid, individuals throughout this paper). Both formulae overestimate the frequency *x*. Eqn. (1) comes from the assumption that the number of new mutations is equal to the number of alleles eliminated by genetic deaths at equilibrium, hence

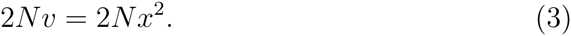

Eqn. (3) should really be

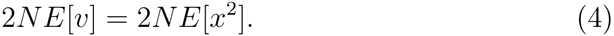

where *E* denotes the expected value. Because (*E[x]*)^2^ ≤ *E*[*x*^2^], Eqn. (1) overestimates *x*. Eqn. (2) is based on the assumption of no recurrent mutation, a mutation becomes extinct before another mutation occurs at that locus. Because subsequent mutations will interact with previous mutations, mutations will be eliminated more quickly and the frequency of deleterious alleles is overestimated.

Since both formulae provide an overestimate of the frequency, the minimum of the two formulae will provide a better estimate of the frequency. These formulae provide the same frequency *x* if 4*πNv* = 1, with 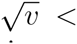 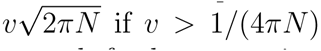 and 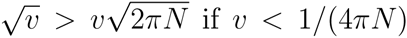. This is expected, for low mutation rates, the nonrecurrent mutation model is reasonable, for high mutation rates, *x*, hence *x*^2^, should have less variation. But *v* < 1/(4*πN*) merely provides that nonrecurrent mutation is a better model than constant mutation frequency, not that it is a good model. Nonrecurrent mutation will be a good model if 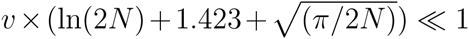 where the second factor is the expected time until extinction of a mutation (Li and Nei, 1972). If *N* > 5, *N* > 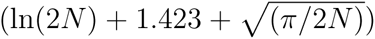 and *v* < 1/(4*πN*) provides 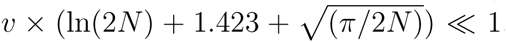. For the remainder of this paper we shall assume *v* is small. In particular, that the equilibrium level of mutations is 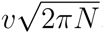 which depends on *N*.

## 3. The mutation rate

The mutation rate is approximately 10^−8^ per base pair per generation (Gao et al., 2015; Harris and Nielsen 2016), which provides approximately 10^−5^ per locus, assuming 1000 base pairs per locus. However, we are interested in the LR mutation rate, which will be less than the total mutation rate. Assuming a human population size of 2*N* = 20,000 (Gao et al. 2015, Harris and Nielsen 2016), a human genome size of 20,000 loci (Spuhler 1948, Ezkurdia et al. 2014), and two LR per diploid individual (Gao et al., 2015), the mutation rate can be calculated from Eqn. (2) using *x* = 1 for the haploid complement producing *v* = .004. Dividing by 20000 to get the mutation rate per locus yields 2 × 10^−7^ which is 2% of the total mutation rate per locus. Gao et al. based their estimate of the mutation rate on LR which manifest death after birth, this does not include deaths that occur in utero, some of which may be due to noncoding regions of DNA. Hence the mutation rate calculated may be low, but such prenatal deaths are more likely to be mitigated by reproductive compensation which will ameliorate their impact. Halligan and Keightly (2003) cite a LR frequency of 1.4 for humans, 1.9 for African clawed frogs and bluefin killifish, 1.6 for the Mexican salamander, 1.4 for zebrafish, a range of 0.5 to 3 for Drosophila, a range of 3 to 6 for Loblolly pine, and higher values for pacific oysters (a range of 8 to 14 (Launey and Hedgecock, 2001)). These values are remarkably similar, since LR frequency should depend on population size.

## 4. Lethal recessive simulations

Simulations were performed of the fate of a single lethal recessive mutation in populations ranging from size 2*N* = 20 to 2*N* = 20000. Each generation the next generation was formed by by sampling with replacement from the present generation; if a homozygous LR individual was formed, another two alleles were sampled so that each generation had 2*N* viable individuals (perhaps carrying one LR gene). The average total number of copies of a LR before it went extinct, the average number of generations until a LR went extinct, the average number of homozygous deaths per mutation, the maximum number of generations until the mutation went extinct, and the average number of generations at which a genetic death (homozygous individual) occurred were recorded. Ten thousand simulations were done thrice for each value of 2*N*, and the middle of the three simulation values are presented in Table 1.

**Table 1:** Results of simulations of single lethal recessive mutations

| 2N | number<br>of copies | time<br>segregating | number<br>of deaths | maximum<br>time | time<br>of deaths |
| --- | --- | --- | --- | --- | --- |
| 20 | 8.50 | 4.34 | .648 | 56 | 7.76 |
| 200 | 26.04 | 6.32 | .538 | 142 | 24.36 |
| 2000 | 77.08 | 8.45 | .477 | 365 | 75.81 |
| 20000 | 246.09 | 10.71 | .491 | 865 | 223.09 |

The first three columns of Table 1 can be obtained from formulae in Li and Nei (1972). The expected number of copies of a mutation is 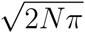 (Li and Nei 1972, Eqn. 7b), this produces the values 7.93, 25.07, 79.27,250.66. The expected time until extinction is ln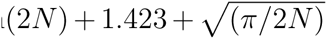 (Li and Nei 1972, p. 675), this produces the values 4.815, 6.847, 9.064, 11.339. The expected number of homozygotes is .5 (Li and Nei 1972, Eqn. 9c), the values in Table 1 should be slightly larger because our model has reproductive compensation for genetic deaths. (Li and Nei used a diffusion approximation, but a martingale argument with a branching process achieves the same result; reproductive compensation changes the martingale to a submartingale.)

## 5. Number of lethal recessive mutations

The expected number of lethal recessive mutations per individual is obtained by multiplying the mutation rate per locus (.0000002) times the number of loci (20000) times the number of copies per mutation 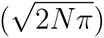 times 2 (for diploid individuals). This produces the values .064, .203, .641, and 2.03 for populations of size 2*N* = 20, 200, 2000, and 20000 (the 2.03 reflects roundoff error, since the mutation rate was calculated assuming the frequency 2 for a population of size 20000.) Note that the results in table 1 are independent of the mutation rate, hence the expected number of LR per individual is proportional to the mutation rate (or the mutation rate per locus times the number of loci). But if the mutation rate is too high, the assumption of nonrecurrent mutation will not be valid.

## 6. Purging load

When a bottleneck occurs, the frequency of LR must be reduced from the equilibrium frequency for the ancestral population size to the equilibrium frequency for the bottleneck population size. We shall calculate this cost three ways: the number of homozygous LR individuals which must be formed (die) to reduce the frequency, the average number of such LR deaths per generation during the period of purging, and the the average number per generation converted to a reduced viability. We illustrate these calculations using the reduction from 2*N* = 20000 to 2*N* = 2000 (motivated by the sizes of the human and Neanderthal populations). The results for this and other transitions are in Table 2. Because the number of LR is proportional to the mutation rate, if the mutation rate is increased, the total number of deaths and average number of deaths in Table 2 will be increased by the same factor.

**Table 2:** The cost of purging lethal recessive mutations

| transition | total<br>deaths | average<br>deaths | average<br>viability |
| --- | --- | --- | --- |
| 20000 → 2000 | 679.5 | 4.48 | .9955 |
| 20000 → 200 | 89.85 | 1.84 | .9816 |
| 20000 → 20 | 9.68 | .6237 | .9376 |
| 2000 → 200 | 21.9 | .4491 | .9955 |
| 2000 → 20 | 2.885 | .1859 | .9814 |
| 200 → 20 | .695 | .04478 | .9955 |

A population of size 2*N* = 20000 has two LR per diploid individual, hence when a bottleneck occurs, that will be the frequency in a population of size 2000. That must be reduced to the equilibrium level for a population of size 2000 which is .641, hence 1000 × (2 — .641) × .5 = 679.5 homozygous deaths must occur. (The number of diploid individuals times the number of LR lost per diploid individual times the number of deaths per LR.) The average time of a LR homozygous death in a population of size 2000 is 75.81, hence most deaths should occur within 2 × 75.81 = 151.62 generations, entailing 679.5/151.62 = 4.48 deaths per generation. If the equilibrium viability is 1, the reduced viability is (1000 — 4.48)/1000 = .9955 for those 151.62 generations.

Although the accumulation of near neutral deleterious alleles is often a primary concern of reduction in population size (Harris and Nielsen 2016), that will occur after LR are purged. The expected time until fixation of a neutral mutation is about 4*N* generations (there is a large variance), which is much larger than the expected time at which purging deaths occur: 40 ≫ 7.76, 400 ≫ 24.36, 4000 ≫ 75.81, and 40000 ≫ 223.09. Therefore, it is appropriate to consider the impact of LR purging in isolation from DM. Using the average time LR segregate as an index of the duration of their impact provides greater inequality, since that is less than the expected time at which purging deaths occur.

One index of the impact of LR is just the number of deaths due to purging. The number of deaths due to purging ranges from 7% of the population size (.695/10 for the transition 200 → 20) to 97% of the population size (9.68/10 for the transition 20000 → 20). In the latter case removing that many individuals from the population would leave less than one individual, and it is reasonable to say the population would go extinct. For the transition 20000 → 2000, two-thirds (679.5/1000) of the individuals would be eliminated.

The deaths could also be converted to a reduction from a mean viability 1 for the population. The result is that over the course of purging for the transition 20000 → 20, .9376^15.52^ = .38, so 38% or four individuals remain. For the transition 200 → 20, .9955^15.52^ = .93 so 93% or 9 of the 10 individuals remain. For the transition 20000 → 2000, .9955^151.62^ = .50 and half the population is lost in the purge.

## 7. Impact on logistic recovery

If a one time event such as a volcano or famine caused the bottleneck, then growth governed by the logistic model (Crow and Kimura, 1970) *dN/dt* = *rN(K - N)/K* where *N* is the population size, *r* is the intrinsic rate of growth, and *K* is the carrying capacity could counter purging deaths due to reduced population size. If the intrinsic rate of growth *r* < .05, the actual growth rate which is reduced by the population size and purging will be less than .05, and the population size will less than treble during a purging period of 20 years (the average time a LR segregates is 10.77 generations for populations of size 10000, less for smaller populations; Leberg and Firmin (2008) note that brief bottlenecks are often only one to three generations). This provides that *(K - N)/K* will remain larger than .7 during a purge for bottlenecks which are 10% of the original population size, even closer to 1 for smaller bottlenecks. Since *(K - N)/K* will be close to 1 during purging, survival will depend on *r* and the cost of purging.

Table 2 suggests a reduction in viability of about .06 if the population size is reduced to .1% of its original size, .02 if the population size is reduced to 1% of its original size, and .005 if the population is reduced to 10% of its original size. Intrinsic rates of growth *r* greater than those values would prevent further population size reduction due to purging. The net effect including when *r* is less than the reduction due to purging can be calculated.

After the LR are purged and the population expands, there will be fewer than the equilibrium number of deaths until the equilibrium level of LR for the larger population size is obtained. The effect of this dearth of deaths increasing viability should be slightly smaller in magnitude than the decrease in viability during the purge because the larger population will have a longer time until equilibrium. There will also be a gradual return to the original population size, as contrasted to a perhaps rapid reduction to a bottleneck which a catastrophe could cause. In summary, if there is logistic population growth, variation in the number of LR deaths will slightly modify but not qualitatively change the nature of the logistic population growth.

## 8. Inbreeding depression and load

Kirkpatrick and Jarne (1999) have said that a bottleneck will always decrease inbreeding depression and the load from recessive mutations will always be increased. It is worth mentioning why this is true in the special case of LR. Inbreeding depression is defined as the loss of fitness in offspring from matings between relatives compared with offspring from random mat-ings within the same population (Kirkpatrick and Jarne, 1999). The fitness of offspring of relatives (especially selfing) will be the same before or immediately after the bottleneck, but random mating will more likely entail selfing after the bottleneck, so this result follows not from increased fitness of inbred matings, but from decreased fitness of random matings. The mutation load is the difference between the fitness of a mutation free population (normalized to 1) and the mean fitness of the population (Crow and Kimura, 1970). The fitness of a mutation free population is the same before or after a bottleneck, but the mean fitness of the population is lower after the bottleneck due to purging (i.e., the purging deaths lower the fitness), so the increased load is due to lower fitness of the bottleneck population.

## 9. Conservation genetics

O’Regan et al. (2002) have noted that although the purges associated with bottlenecks threaten populations with extinction, if the purged population has not reached the equilibrium level of LR for its recovered large size, it will not be subject to as much purging during the next bottleneck. This is true, but the equilibrium level of LR is reached in only a few times the average number of generations segregating, and that time only increases as the logarithm of the population size. Therefore the time to restore the enlarged population to equilibrium is not much larger than the time to complete a purge, and there will generally not be a reduced level of LR when the next bottleneck occurs. However, if the small population size is maintained, a subsequent bottleneck will be less serious as noted by Leberg and Firmin (2007) (The cost of going from 200 to 20 is less than the cost of going from 2000 to 20, and if the reduction is done in two steps, the viability reduction is less). But in this circumstance, DM may accumulate due to drift. Purging is not a useful strategy for conservation genetics.

Indeed, small population size should be avoided when managing and restoring populations because deleterious mutations may be fixed due to drift. But as noted by Leberg and Firmin (2007), the time required for bottlenecks to eliminate LR is briefer than the time necessary for DM to accumulate, hence a brief size reduction to eliminate LR should not be a problem. This could be important for the conservation genetics strategy of breeding individuals from large populations who will not have DM fixed at several loci into endangered populations. A small subpopulation of the large population could be formed and maintained for a few generations to purge the LR without allowing time for DM to accumulate, so that LR are not introduced into the endangered population. Purging LR in a captive population could prevent random (e.g., predator) deaths of viable individuals, enhancing the probability of the subpopulation surviving the purge.

## 10. Discussion

The number of genetic deaths due to lethal recessive mutations is equal to one-half the mutation rate (each death removes two LR). This is independent of population size, but the equilibrium level of LR depends on the population size. When the population size changes, the frequency of LR does not immediately change, and this causes more or fewer LR deaths than half the mutation rate until equilibrium is achieved for the population size.

But equilibrium is quickly attained. The purging deaths in a bottleneck will indeed exacerbate the conditions which caused the bottleneck, but if the conditions which caused the bottleneck are no longer present, a logistic return to the previous population size will overcome the cost of purging. The benefits of purging a population of LR are lost almost as quickly as the LR were purged.

LR mutations do not impact the accumulation of DM. LR mutations quickly achieve equilibrium for any population size, DM slowly drift to fixation or extinction. The impact of DM occurs long after a purge is over.

